# Brainstem cholinergic cell loss causes slowly-developing motor impairments

**DOI:** 10.64898/2026.10.16.759730

**Authors:** K Udobi, V Aleman, VJ Lewitus, M Fallah, BJ Bernstein, G Zhang, M Ezeiza Ortega, C Konja, S Grossen, P Forcelli, N Neckel, RC Evans

**Author notes:** Contributed equally to this work (K Udobi, V Aleman, VJ Lewitus). The authors declare no conflicts of interest.

## Abstract

Pedunculopontine nucleus (PPN) cholinergic degeneration is a feature of Parkinson’s disease (PD), but it is difficult to isolate the symptoms and circuit alterations caused by cholinergic cell loss in PD, where multiple nuclei are simultaneously degenerating. Here we develop a novel mouse model of brainstem cholinergic cell loss to define the timeline of symptom development and map physiological adaptations within the brainstem. Mild motor symptoms, including balance and motor learning deficits reminiscent of early-stage PD, developed after cholinergic lesion with an unexpected long delay. *Ex vivo* whole-cell recordings showed that cholinergic cell loss increased excitatory synaptic drive onto glutamatergic PPN neurons and increased their intrinsic ability to generate sustained action potential output. In parallel, fiber photometry experiments revealed a slight increase in glutamatergic sensory responsiveness that closely tracked the delayed onset of behavioral symptoms. Together, these results reveal that brainstem cholinergic cell loss triggers the reorganization of motor-related brainstem circuits, providing critical insight into the consequences of non-dopaminergic degeneration in PD.

## Introduction

While the substantia nigra pars compacta (SNc) degeneration is the hallmark of Parkinson’s disease (PD) and SNc lesions recapitulate many cardinal PD symptoms, other midbrain structures also degenerate in PD. Specifically, the pedunculopontine nucleus (PPN), a brainstem structure implicated in gait, posture, and balance control, degenerates in PD ^1–6^. The PPN contains cholinergic, glutamatergic, and GABAergic neurons ^7^ that differentially modulate motor activity ^8–14^. PPN cholinergic neurons are particularly susceptible to pathological factors ^15,16^ and most heavily degenerate in PD ^1,5^. However, it has been difficult to determine which symptoms and circuit alterations result from cholinergic cell loss in PD because multiple neuronal populations degenerate simultaneously. Because some PD symptoms can be resistant to dopamine replacement therapy ^17,18^, but may respond to PPN deep brain stimulation ^19–23^, it is important to understand how degeneration in the PPN contributes to the motor and non-motor aspects of PD.

Previous studies have made important findings using toxin-based brainstem cholinergic lesions primarily in male rats. These studies report mild PD-like symptoms including altered gait ^4,24^, impaired startle response ^25^, and motor skill deficits ^26–28^, but no change in reward seeking ^29^. However, creating and characterizing a mouse model of brainstem cholinergic cell loss is critical for fully integrating the genetic tools available in mice to comprehensively dissect the symptoms, cellular properties, and circuit alterations related to cholinergic cell loss in PD. Interestingly, while the cholinergic neurons are the ones that degenerate in PD, it is the glutamatergic PPN neurons that have been primarily implicated in motor control ^9,11,13,30,10,31^. Because glutamatergic and cholinergic neurons are intermixed within the PPN ^7^, it is important to determine whether PPN cholinergic cell loss alters the cellular properties and circuit connectivity of neighboring glutamatergic neurons.

Here, we develop a novel mouse model of PD-related brainstem cholinergic cell loss and detail the time course of motor and non-motor symptom development. We use *ex vivo* whole-cell electrophysiology to reveal intrinsic and synaptic physiological changes in opto-tagged glutamatergic PPN neurons after cholinergic cell loss. Using fiber photometry, we measure the activity of brainstem glutamatergic neurons in control and brainstem cholinergic cell loss conditions. We find that brainstem cholinergic cell loss results in symptoms similar to those seen in dopamine-mediated early-stage and progressive PD models. Interestingly, symptoms develop weeks after the cholinergic lesion and are accompanied by a mild increase in glutamatergic responsiveness to sensory stimuli developing with a similar time course. During this same time-frame, excitatory synaptic input onto PPN glutamatergic neurons is strengthened while intrinsic excitability is reduced in cholinergic cell loss conditions. These findings support the idea that changes in cellular and circuit plasticity slowly develop in response to brainstem cholinergic cell loss.

## Results

### Caspase-dependent cholinergic cell loss along PPN rostral-caudal axis

To selectively lesion cholinergic neurons in the pedunculopontine nucleus (PPN), we injected an AAV containing a Cre-dependent caspase bilaterally into the PPN of ChAT-Cre/Ai9-tdTomato mice. Wild type (WT) controls were injected with the Cre-dependent caspase as a virus control, and ChAT-Cre/Ai9-tdTomato mice were injected with a Cre-dependent GFP virus as a genotype control (Figure 1A). Using CUBIC clearing ^32^, confocal imaging, and cell counting with a custom ImageJ macro, we found a reduction in cholinergic neurons in the lesion condition (Figure 1B). Directly comparing the number of tdTomato-positive neurons in lesion and control ChAT-Cre/Ai9-tdTomato mice, we see clear reduction across the rostral-caudal extent of the PPN (group effect: F_(1,28)_ = 27.03, p < 0.0001, linear mixed-effects model, Figure 1C). The presence of PPN cholinergic neurons in WT control mice was verified with antibody staining (*data not shown*). Because staining vs genetic labeling is not directly comparable, we did not directly compare antibody-based cell counts to tdTomato-based cell counts.

**Figure 1.**
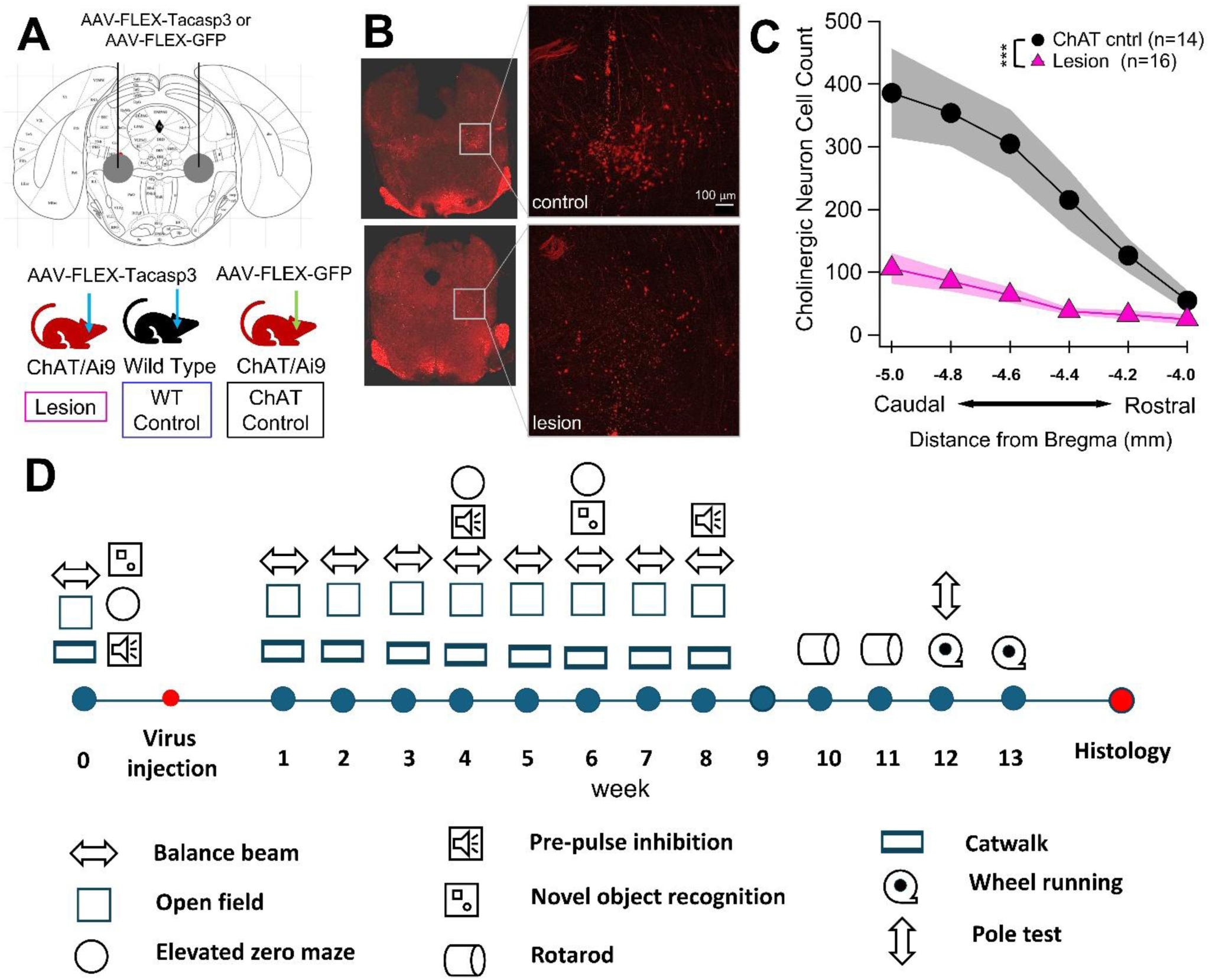
Lesion of cholinergic PPN neurons. **A.** Bilateral injections into ChAT/Ai9-tdTomato and WT littermates. **B.** Confocal image of PPN in control (top) and lesion (bottom) mouse. **C.** In the lesion condition, cholinergic neuron cell loss spans the rostral-caudal extent of the PPN. **D.** Timeline for behavior experiments. *** p<0.0001

### PPN cholinergic cell loss induces motor symptoms, impairing motor skill recall and balance

The PPN is implicated in balance and postural stability deficits seen in PD. Parkinson’s patients with falling as a main symptom have more extensive cholinergic PPN degeneration than PD patients without falling as a main symptom ^4,6^. Previous work with toxin-based PPN cholinergic lesions show postural stability deficits in monkeys ^4^, but no impairments in traversing a narrow beam in male rats ^27^. Interestingly, a recent study actually shows balance improvements during chemogenetic inhibition of PPN cholinergic input to the cerebellum in rats ^33^. To determine whether balance was disrupted in our brainstem cholinergic cell loss mouse model, we assessed the time to cross parallel 3 mm-wide balance beams weekly for 8 weeks post-injection (Figure 1D). In order to test balance skill rather than motor skill learning, mice were trained for 5 days prior to viral injection. Only mice that met the performance criteria (crossing within 120 seconds on the last day) were included in balance beam behavioral analysis and data were normalized to baseline performance for each mouse. Importantly, there were no differences in baseline balance beam performance across groups (WT: 35.86 ± 6.69 s, n = 19; ChAT: 25.29 ± 7.69 s, n = 14; lesion: 32.86 ± 7.45 s, n = 15; F_(2, 42)_ = 0.55, p = 0.58 one-way ANOVA). After viral injection, mice with brainstem cholinergic cell loss showed a slowly-emerging balance deficit, taking significantly longer than the control groups to cross the balance beams at 6 and 7 weeks post-injection. There was a significant group × week interaction (F_(14,315)_ = 1.81, p = 0.037, linear mixed-effects model) and post-hoc tests showed significantly longer times to cross in lesion conditions compared to both control groups at 6- and 7-weeks post-lesion (Week 6 WT: 132 ± 29.3% n=19, ChAT: 115 ± 12.4% n=14, lesion: 335.857 ± 111.16% n=15 Les vs WT t_(315)_=2.94, p = 0.0068; Les vs ChAT t_(315)_=2.96, p= 0.0064, Week 7 lesion: 321.3 ± 128.23%, WT: 129.34 ± 36.6%, ChAT: 114.8 ± 16.4% Les vs WT t_(315)_=2.77, p = 0.012, Les vs ChAT t_(315)_=2.77, p=0.011, Dunnett’s Figure 2A), but not at 8-weeks post-lesion (Week 8 lesion: 222.6 ± 103.02%, WT: 140.86 ± 36.6%, ChAT: 187.4 ± 61.1% Les vs WT t_(315)_=1.18, p = 0.24, Les vs ChAT t_(315)_=0.47, p=0.64, Dunnett’s), reflecting delayed emergence of balance deficits.

**Figure 2.**
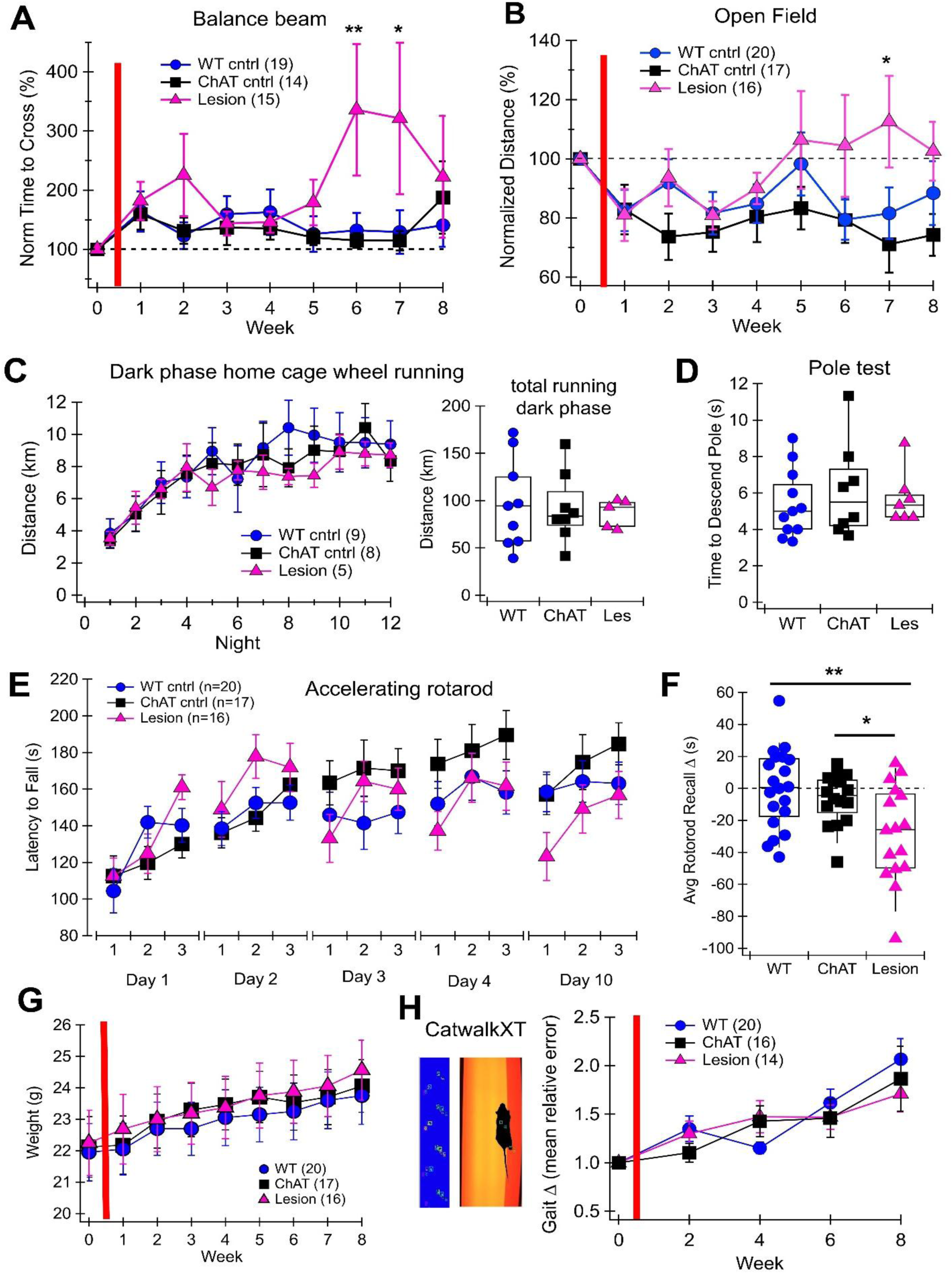
Cholinergic cell loss reduces motor skill. **A**. Time to cross a 3mm wide pair of balance beams. Red line indicates time of lesion or control injection. Lesion mice take longer to cross the beam 6- & 7-weeks post-injection. **B.** Distance traveled in 10 minutes of open field exploration tested weekly. Lesioned mice show increased exploration 7 weeks post-injection. **C.** Home cage running amount or pattern does not differ in lesion vs control mice. **D.** Time to turn and descend pole (average of 3 trials on testing day) **E.** Latency to fall on the accelerating rotarod. Lesioned mice show a deficit in motor skill retention from one day to the next, but show motor skill learning within each daily session. **F.** Lesioned mice show significant deficits in motor skill recall on the rotarod. First trial of day minus last trial of previous day (averaged per mouse across all days). **G.** Mouse weight steadily increased over experimental time period, but did not differ across groups. **H.** Gait analysis shows no significant gait abnormalities across groups. **p<0.01 *p<0.05

To determine whether there were any gross motor deficits in our brainstem cholinergic cell loss mouse model, we allowed the mice to explore a plexiglass open field for 10 minutes starting the week prior to viral injection and continuing weekly for 8 weeks post-injection. No baseline locomotion criteria were applied and data were normalized to the pre-lesion distance traveled for each mouse. We found trend effect of week and group on distance traveled in the open field (Week: F_(7,350)_ = 1.87, p = 0.073, Group: F_(2,50)_ = 2.34, p = 0.107, linear mixed-effects model, Figure 2B). Testing slice effects showed that lesion mice traveled significantly longer distances at 7-weeks post-injection (Week 7 WT: 81.58 ± 8.7%, n = 20; ChAT: 71.14 ± 9.6% n=17, lesion: 112.56 ± 15.52% n = 16 Les vs WT t_(350)_=2.43 p=0.029; Les vs ChAT t_(350)_=3.04, p=0.005 Dunnett’s) and trended toward longer distances at 6-weeks post-injection (Week 6: Les vs WT t_(350)_=1.95, p=0.09; Les vs ChAT t_(350)_=1.84, p=0.12). These data indicate that locomotor hyperactivity progressively emerges during the post-lesion period. This finding is surprising because PD is characterized by reduced locomotion. However, a hyperactive phenotype has been observed during the earliest stages of disease development in progressive PD animal models and may relate to a period of dopaminergic hyperactivity ^34,35^. Thus, an increase in locomotion could represent an early-stage change in the dopaminergic system occurring in response to cholinergic cell loss.

The PPN has been implicated in exploratory locomotion, but not escape-related movements like running or darting ^8^. Therefore, we evaluated home-cage wheel running to determine whether home-cage hyperactivity appears in parallel with increased exploratory locomotion in the open field (Figure 2B). Because home-cage running strongly affects PPN cholinergic neuron activity ^36,37^, we added tracking running wheels to the home cage after all other behavioral tests were complete. We found that the total amount of home cage running summed across all 12 nights did not differ across groups (total dark phase running lesion: 86.73 ± 6.68, n = 5; WT: 97.23 ± 15.73, n = 9 ChAT: 91.94 ± 12.92, n = 8; one-way ANOVA: F_(2,21)_ = 0.124, p = 0.88, Figure 2C, *right*). This indicates that the hyperactivity phenotype in this model is specific to exploration of an open field environment and is not reflected in higher levels of home-cage wheel running.

An important symptom in PD is the impairment of motor skill learning ^38–40^. Multiple early-stage PD animal models that involve partial dopaminergic cell loss show motor skill learning deficits in the absence of gross locomotor impairments ^41–43^, and an toxin-based PPN lesions can impair accelerating rotarod performance in male rats ^26^. To determine whether motor skill learning was impaired by selective PPN cholinergic cell loss, we ran the accelerating rotarod assay. This test was run only once per mouse during weeks 10-11 post-injection. On the accelerating rotarod, mice with brainstem cholinergic cell loss showed motor skill loss from one training session to the next (Figure 2E). Motor skill recall was calculated as the difference between the latency to fall in the first trial of the session and the last trial from the previous session and averaged per mouse over all days (Figure 2F). Mice with cholinergic cell loss had significantly impaired motor skill recall compared to both control groups (lesion: -28.02 ± 7.52, n = 16; WT: -0.325 ±5.52, n = 20 ChAT: -6.48 ± 3.94, n = 16; one-way ANOVA: F_(51)_ = 6.13, p = 0.004; lesion vs WT p=0.0026; lesion vs ChAT p=0.029, post-hoc Dunnett’s test). Importantly, these motor recall deficits occurred in the absence of group-dependent differences in weight (Figure 2G) or severe disruptions to motor coordination, as measured by the pole test (Figure 2D; lesion: 5.7 ± 0.55, n = 7; WT: 5.42 ± 0.57, n = 11 ChAT: 6.13 ± 0.91, n = 8; Kruskal-Wallis: H = 0.49, p = 0.76). These results show that PPN cholinergic lesions impair recall of previously acquired motor learning, but do not disrupt within-session motor skill learning. This finding contrasts with the complete lack of motor skill learning observed in partial dopamine depletion animal models ^41,43^, but is consistent with a human study showing PD patients have relatively normal short-term motor skill learning, but impaired motor skill recall 24 hours later ^40^. These data indicate that PPN cholinergic neurons have a specific role in motor skill memory consolidation and recall.

Because the PPN is implicated in freezing of gait and falling in PD patients ^4,6,44^, we evaluated the effects of PPN cholinergic cell loss on mouse gait using the Noldus CatWalkXT before and after viral injections. Gait parameters such as stride length, limb phasing, and base of support were measured and consolidated into mean relative error for each mouse ^45^. This error value was normalized to gait error prior to viral injection (week 0) for each mouse. While all error values increased over weeks (F_(3,149)_=8.94, p<0.001, linear mixed-effects model), there were no significant difference across groups at any time point (week 2: F_(2,149)_=0.44, p=0.64; Week 4: F_(2,149)_=1.3 p=0.28; Week 6: F_(2,149)_=0.1 p=0.91; Week 8: F_(2,149)_=0.8 p=0.45, linear mixed-effects model, Figure 2H). Therefore, these data show that brainstem cholinergic cell loss does not significantly impair gait on its own.

### PPN cholinergic cell loss does not impair several non-motor behaviors

Because anxiety is a common and debilitating non-motor symptom in PD ^46^, we tested anxiety-like behaviors in the brainstem cholinergic cell loss model. We evaluated the time mice spent in the center area of an open field, and the time mice spent in the open vs closed arms on an elevated zero maze. Time spent in center or open arm was normalized to the pre-surgery value for each mouse to evaluate any changes over time. There was no difference in the normalized time in center across groups or weeks and no interaction effect (week F_(7,350)_=1.65, p=0.12; group F_(2,50)_=0.79, p=0.46; group x week F_(14,350)_=0.60, p=0.86; linear mixed-effects model, Figure 3A). In addition, time spent in open sections of the elevated zero maze was normalized to pre-surgery values and did not differ by group or week (week F_(2,102)_=2.39, p=0.096; group F_(2,51)_=0.74, p=0.64; group x week F_(4,102)_=0.53, p=0.71; linear mixed-effects model Figure 3B). Together these data show that brainstem cholinergic cell loss does not affect anxiety-like behaviors.

**Figure 3.**
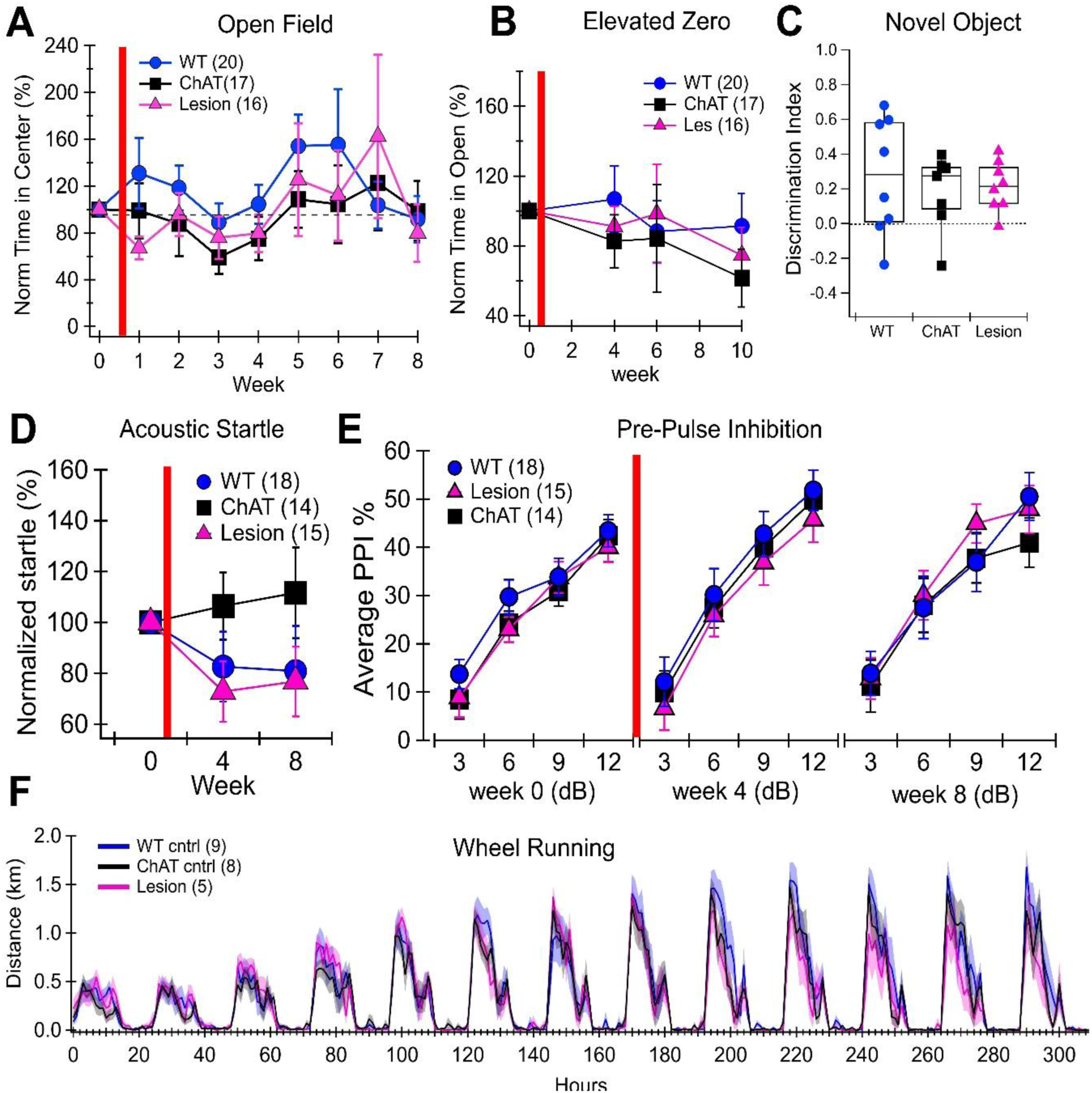
Brainstem cholinergic cell loss does not impair several non-motor behaviors.. **A.** Time in center region of open field does not differ by week or group. **B.** No effect on time spent in open arms of the elevated zero maze. **C.** Novel object recognition memory is not impaired. **D.** Acoustic startle response did not significantly differ by group. **E.** Pre-pulse inhibition (PPI) was not impaired by cholinergic lesion. **F.** Circadian rhythm measured with home-cage wheel running was not altered by cholinergic cell loss.

Because healthy mice adapt to an open field and move less with repeated exposure ^47^, the increased open field locomotion we observed in lesioned mice (Figure 2B) may reflect impaired habituation to the familiar environment. We hypothesized that mice with cholinergic cell loss may not recognize the open field box as familiar. To determine if recognition memory was impaired in the brainstem cholinergic cell loss condition, we used the novel object recognition task 6 weeks post-injection. Mice were exposed to a familiar and novel object and their ability to discriminate between the two was calculated as discrimination index = (time spent exploring novel – time spent exploring familiar) / total exploration time. Discrimination index did not differ across groups (Lesion 0.21 ± 0.05, n = 8; WT 0.27 ± 0.12, n = 8; ChAT 0.18 ± 0.08, n = 7; one-way ANOVA F_(22)_ = 0.29, p = 75, Figure 3C), indicating that object recognition memory is intact after brainstem cholinergic cell loss. This is in contrast to a previous study which found object recognition deficits using an ibotenic acid PPN lesion model (which is not selective for cholinergic neurons) in rats ^48^. It is particularly interesting that we find impairments in motor skill recall (Figure 3E) but not novel object recognition. This could reflect the recall timeframe, as novel object recall was tested 1 hour after exposure to the familiar objects, but motor recall was tested on the scale of days and weeks. Together, these data indicate that brainstem cholinergic neurons are critical for specific types of memory or specific timeframes for recall.

The PPN has been implicated in sensory gating and acoustic startle response in healthy and parkinsonian conditions ^49–52^. Previous toxin-based PPN cholinergic lesions in male rats show acoustic startle deficits ^25^, but ChAT knockout conditions can enhance pre-pulse inhibition (PPI) ^53^. Optogenetic stimulation of cholinergic PPN neurons enhances acoustic startle responses ^54^, while optogenetic stimulation of glutamatergic PPN neurons can reduce it at short inter-stimulus-intervals ^55^. To determine whether acoustic startle or sensory gating was altered in our brainstem cholinergic cell loss model, we ran the PPI assay before viral injections and at two time points after (Figure 1D). Startle habituation was assessed by acoustic startle response immediately before and after the PPI assays. If startle amplitudes after PPI tests were lower than 40% of baseline startle amplitude, data were excluded. Data were normalized to pre-surgery startle amplitude for each mouse. We found that PPN cholinergic lesions did not significantly impair acoustic startle response (week F_(1,44)_=0.38, p=0.54; group F_(2,44)_=1.41, p=0.26; group x week F_(2,44)_=0.31, p=0.74; linear mixed-effects model, Figure 3D). As expected, pre-pulse stimulation paired with startle stimuli reduced startle response, with lower amplitude pre-pulses (e.g. 3 dB above baseline) being less effective than higher amplitude pre-pulses (e.g. 12 dB above baseline) (Figure 3E). However, there was no effect of cholinergic cell loss on pre-pulse inhibition and no interaction between group and PPI dB level or week (group F_(2,44)_=0.21, p=0.81; group x week F_(2,44)_=0.96, p=0.39; group x week x PPI level F_(6,132)_=0.83, p=0.55; linear mixed-effects model). These data show that brainstem cholinergic cell loss does not significantly impact acoustic startle or sensory gating.

The PPN is implicated in sleep-wake regulation ^56^. To determine whether circadian rhythm was altered with cholinergic cell loss, we tracked the light/dark wheel running patterns. Cholinergic cell loss did not alter home cage activity patterns (Figure 3F). Therefore, our data show that none of non-motor symptoms tested were altered by PPN cholinergic cell loss.

### PPN glutamatergic neurons have increased excitatory synaptic input after cholinergic cell loss

While it is the cholinergic PPN neurons that primarily degenerate in PD ^2,57^, it is the glutamatergic PPN neurons that have been most clearly implicated in motor control ^9,11,13,10,31^. Because glutamatergic and cholinergic neurons are intermixed within the PPN ^7^, we hypothesized that the loss of cholinergic PPN neurons would alter the electrophysiological properties of the remaining PPN glutamatergic neurons. To test this hypothesis, we injected the cre-dependent caspase (or vehicle control) with a CamKIIa-promoted channelrhodopsin (ChR2) into the PPN, allowing for *ex vivo* electrophysiological recordings of opto-tagged glutamatergic PPN neurons (CaMKIIa-positive) in lesion and control conditions. Coronal brain slices containing the PPN were collected and glutamatergic PPN neurons were recorded 5-8 weeks after injection. ChR2-positive status was confirmed by the presence of sustained inward currents in response to blue (470nm) light stimulation (Figure 4A). Spontaneous excitatory post-synaptic currents (sEPSCs) were recorded from opto-tagged PPN neurons in whole-cell voltage clamp, holding at -70mV (Figure 4B). We found that PPN glutamatergic neurons receive larger amplitude sEPSCs in the lesion condition than in the control condition (Lesion mdn = -23.81 pA, IQR = 12.62 pA, n = 24; Control mdn = -15.94 pA, IQR = 6.66 pA, n = 16; Wilcoxon Mann-Whitney U_(24,16)_ = 110, p = 0.023, Figure 4C). The sEPSC frequency did not differ between lesion and control conditions (Lesion mdn = 2.16 Hz, IQR = 2.87 Hz, n = 24; Control mdn = -1.92 Hz, IQR = 1.77 Hz, n = 16; Wilcoxon Mann-Whitney U_(24,16)_ = 231, p = 0.29, Figure 4D). The sEPSC kinetics also did not differ in either decay tau (Lesion 2.1 ± 0.12 ms, n = 24; Control 2.6 ± 0.16 ms, n = 16; 2-tailed T-test T_(23,15)_ = -1.57, p = 0.13, Figure 4E) or 10-90% rise time (Lesion 0.72 ± 0.03 ms, n = 24; Control 0.76 ± 0.04 ms, n = 16; T-test T_(23,15)_ = -0.86, p = 0.39, Figure 4F). Together, these data show that cholinergic cell loss results in a strengthened excitatory drive onto glutamatergic PPN neurons without changing probability of pre-synaptic release or excitatory current characteristics.

**Figure 4.**
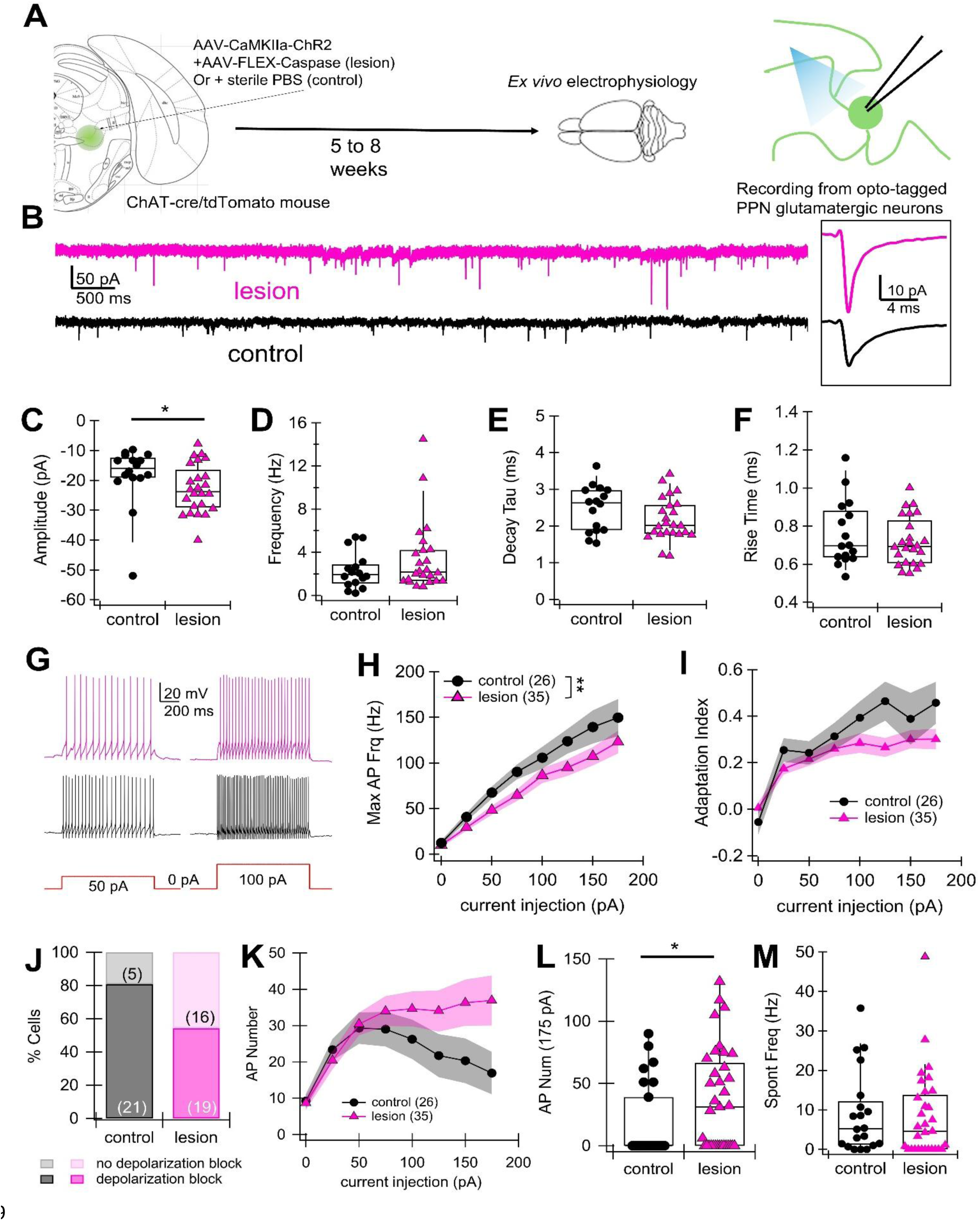
Excitatory input onto glutamatergic PPN neurons is strengthened by cholinergic cell loss.. **A.** Schematic of ex vivo electrophysiology experiments with opto-tagging. **B.** Example raw spontaneous excitatory post-synaptic current (sEPSC) voltage clamp recording from opto-tagged PPN glutamatergic neurons in lesion (pink) and control (black) conditions. Inset = averages sEPSC for each cell. **C.** Average sEPSC amplitude is significantly larger in lesions vs control conditions. **D.** Average sEPSC frequency. **E.** Average decay tau **F.** 10-90% rise time **G.** Example current clamp recording with 50 and 100 pA current injections for lesion (pink) and control (black) conditions. **H.** Maximum instantaneous frequency recorded at each current depolarization step. **I.** Adaptation index at each current step. **J.** Percent of recorded glutamatergic PPN neurons that went into depolarization block at 175 pA or lower current injection. **K.** Total action potential number during 1 second current injection. **L.** Action potential number during 175pA current injection. **M.** Spontaneous action potential firing frequency. **p<0.01 *p<0.05

To evaluate the intrinsic excitability of PPN glutamatergic neurons, we injected depolarizing current steps through the patch pipette in current clamp mode (Figure 4G). Measuring action potential response at each current step, we found that there was a significant effect of group on the maximum firing rate (F_(1,66.8)_ = 8.38, p = 0.0051, linear mixed-effects model, Figure 4H). These data show that cholinergic cell loss reduces glutamatergic PPN intrinsic excitability. To measure the ability of PPN glutamatergic neurons to maintain a sustained output in response to current input, we measured the spike frequency adaptation at each current step, calculated as 1-(final frequency/initial frequency). There was no significant effect of group on adaptation index (F_(1,65)_ = 1.32, p = 0.25, linear mixed-effects model, Figure 4I). However, there is a non-significant trend toward a lower percentage of PPN glutamatergic neurons entering depolarization block during these current injections in lesion (54%, 19/35) vs control (80.7% 21/26) condition (p=0.055, two-sided Fisher’s Exact Test, Figure 4J). To further investigate the ability of these neurons to sustain firing, we counted the total action potential number during each current injection. This increases with increasing depolarization, but decreases if a neuron enters depolarization block. PPN glutamatergic neurons from the lesion condition maintain firing frequency at higher current injections, while neurons in the control conditions decrease firing due to entering depolarization block (Figure 4K). At the highest current injection, 175 pA, action potential number was significantly larger in lesion vs control conditions (Lesion mdn = 31, IQR = 66.5, n = 35; Control mdn = 0, IQR = 39, n = 26; Wilcoxon Mann-Whitney U_(26,35)_ = 320, p = 0.03, Figure 4L). These data show that PPN glutamatergic neurons are able to maintain sustained firing more effectively in the absence of cholinergic neurons. Interestingly, spontaneous action potential firing frequency was not altered by cholinergic cell loss (Lesion mdn = 4.6 Hz, IQR = 13.81 Hz, n = 31; Control mdn = 5.24 Hz, IQR = 10.96 Hz, n = 20; Wilcoxon Mann-Whitney U_(20,31)_ = 343.5, p = 0.52, Figure 4M), indicating that intrinsic excitability and baseline spontaneous activity cells are differentially modulated by PPN cholinergic neurons. These data show that the maximal firing of PPN glutamatergic neurons is reduced, but ability to sustain constant firing is increased, when cholinergic PPN neurons are lesioned. Together, the combination of increased synaptic excitation (Figure 4B&C) and reduced intrinsic excitability (Figure 4H) indicates that cholinergic cell loss modifies glutamatergic brainstem cellular physiology and circuit connectivity.

### PPN glutamatergic neurons respond to visual stimuli with and without cholinergic cell loss

To determine whether cholinergic cell loss altered PPN glutamatergic activity in the intact brain, we used fiber photometry to measure PPN activity with and without cholinergic cell loss over a 6-week time period. PPN glutamatergic neurons respond to sensory stimuli more strongly than other PPN neural populations ^58^. Therefore, we hypothesized that PPN cholinergic cell loss would influence PPN glutamatergic neural responses to sensory input. To evaluate PPN glutamatergic population activity, we injected a CaMKIIa-promoted GCaMP8f with either PBS (control) or the cre-dependent caspase (lesion) and implanted an optical fiber in the PPN of ChAT-cre/Ai9-tdTomato mice (Figure 5A). Three weeks following viral injections, we performed fiber photometry to record bulk calcium dynamics from glutamatergic PPN neurons in both control and caspase-lesioned cohorts during visual stimuli. When room lights were turned on, glutamatergic PPN neurons showed a large transient calcium increase (Figure 5B, *left)*. When room lights were turned off, the glutamatergic PPN neurons showed a much smaller transient calcium increase (Figure 5B, *right*). This shows that PPN glutamatergic neurons do not communicate information about whether the light is currently on or off, but rather send transient signals communicating a change in sensory input.

**Figure 5.**
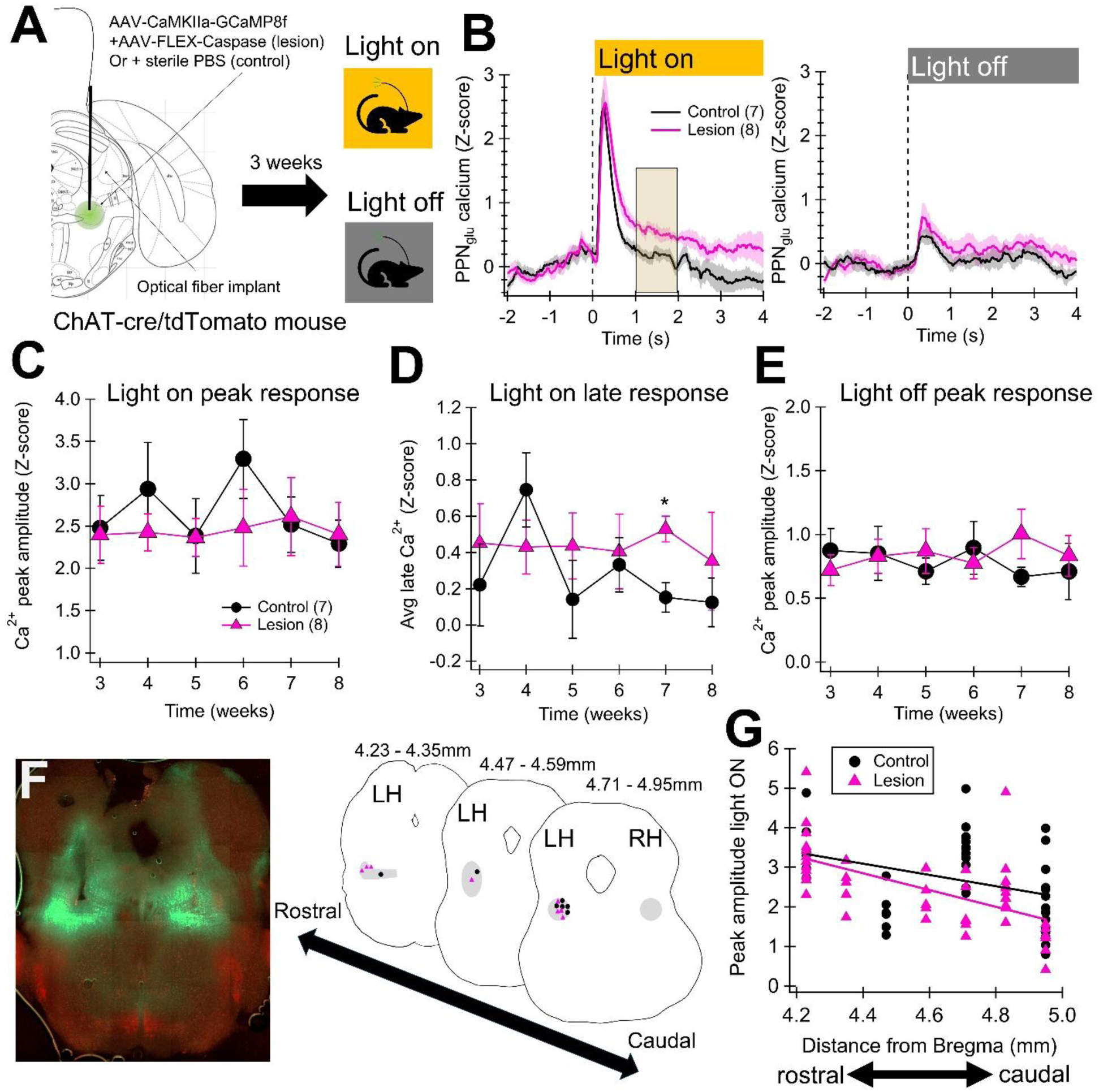
Glutamatergic PPN response to visual stimuli is minimally altered by cholinergic cell loss.. **A.** Experimental setup. GCaMP8f expressed in CaMKIIa-expressing (putatively glutamatergic) PPN neurons and an optical fiber implanted over the PPN in lesion and control conditions. **B.** Example traces showing glutamatergic PPN calcium activity when the room lights turn on (*left*) and off (*right*). Traces are from 7 weeks post-injection time point. **C.** Peak amplitude of glutamatergic PPN calcium response to light turning on is stable over weeks and does not differ between control and lesion conditions. **D.** Average late glutamatergic PPN calcium response measured at shaded area in B (1-2 seconds after light turns on), is larger when cholinergic PPN neurons are lesioned, but only at 7 weeks post-injection. **E.** PPN glutamatergic neurons show a small response to room lights turning off that is stable over weeks and does not differ between control and lesion conditions. **F.** Example image and schematic of implant locations for control (black) and lesion (pink) conditions. **G.** Peak glutamatergic PPN calcium amplitude correlates with rostral-caudal implant location within the PPN in both control and lesion conditions. *p<0.05

The amplitude of the primary PPN glutamatergic response to the light turning on and off were both stable over a period of 6 weeks and did not differ between the cholinergic lesion condition and control (Figure 5C&E). However, there was an increase in the calcium response 1-2 seconds after the light turned on in the cholinergic lesion condition only at 7-weeks post-injection (T_(12.41)_=3.45 p=0.027, T-test Bonferroni correction, Figure 5D). This shows that the glutamatergic PPN neurons took longer to return to baseline activity levels in the absence of cholinergic neurons. It is interesting that this alteration in glutamatergic PPN response to sensory stimuli occurred only at 7 weeks post-injection because that time point showed the strongest behavioral differences in a separate set of mice (Figure 2A&B). Together, these data indicate that both circuit and behavior alterations preferentially occur seven weeks after caspase injection.

The PPN has a distinct topography and exhibits functional and connectivity differences across its rostral-caudal axis ^59–61,30,62,63^. After fiber photometry experiments, brains were sliced and cleared for post-hoc imaging. We identified the location of each implant and recorded the distance from bregma of each location to compare responses across the rostral-caudal axis of the PPN (Figure 5F). We found a significant effect of the distance from bregma on the peak glutamatergic response to the light turning on (β=-2.12 F_(1,75)_=5.04 p=0.028, nested (6 recordings per mouse) linear mixed-effects model, Figure 5G). This effect did not differ between control and lesion conditions (F_(1,75)_=0.19 p=0.66). These data show that glutamatergic neurons in the rostral PPN respond more strongly to visual stimuli than those in the caudal PPN and that this occurs both in the presence and absence of cholinergic PPN neurons.

### PPN glutamatergic neurons respond to auditory and vestibular stimuli with and without cholinergic cell loss

The PPN has been implicated in auditory sensory gating and acoustic startle response ^51,52,54,55^. To evaluate the effect of cholinergic cell loss on glutamatergic PPN response to auditory stimulation, we recorded glutamatergic activity using the same fiber photometry recording set-up as described above. For auditory stimulation, mice were placed in an open field in the presence of 70 dB background white noise and exposed to a 500 ms, 82 dB, 2800-3800 Hz tone given 90 seconds apart following a 150 second baseline (Figure 6A). PPN glutamatergic population response at the time of the tone was recorded 4 times and averaged for each mouse. The amplitude of the glutamatergic PPN calcium change was calculated as the difference between the baseline Z-score prior to the tone and the peak immediately after the tone. There was no difference in calcium response amplitude between the control and lesion conditions (Lesion 1.86 ± 0.50 SD, n = 7; Control 1.12 ± 0.32 SD, n = 8; 2-tailed T-test T_(7,6)_ = -1.24, p = 0.24, Figure 6B). These data show that cholinergic cell loss does not significantly alter the glutamatergic PPN response to auditory stimulation.

**Figure 6.**
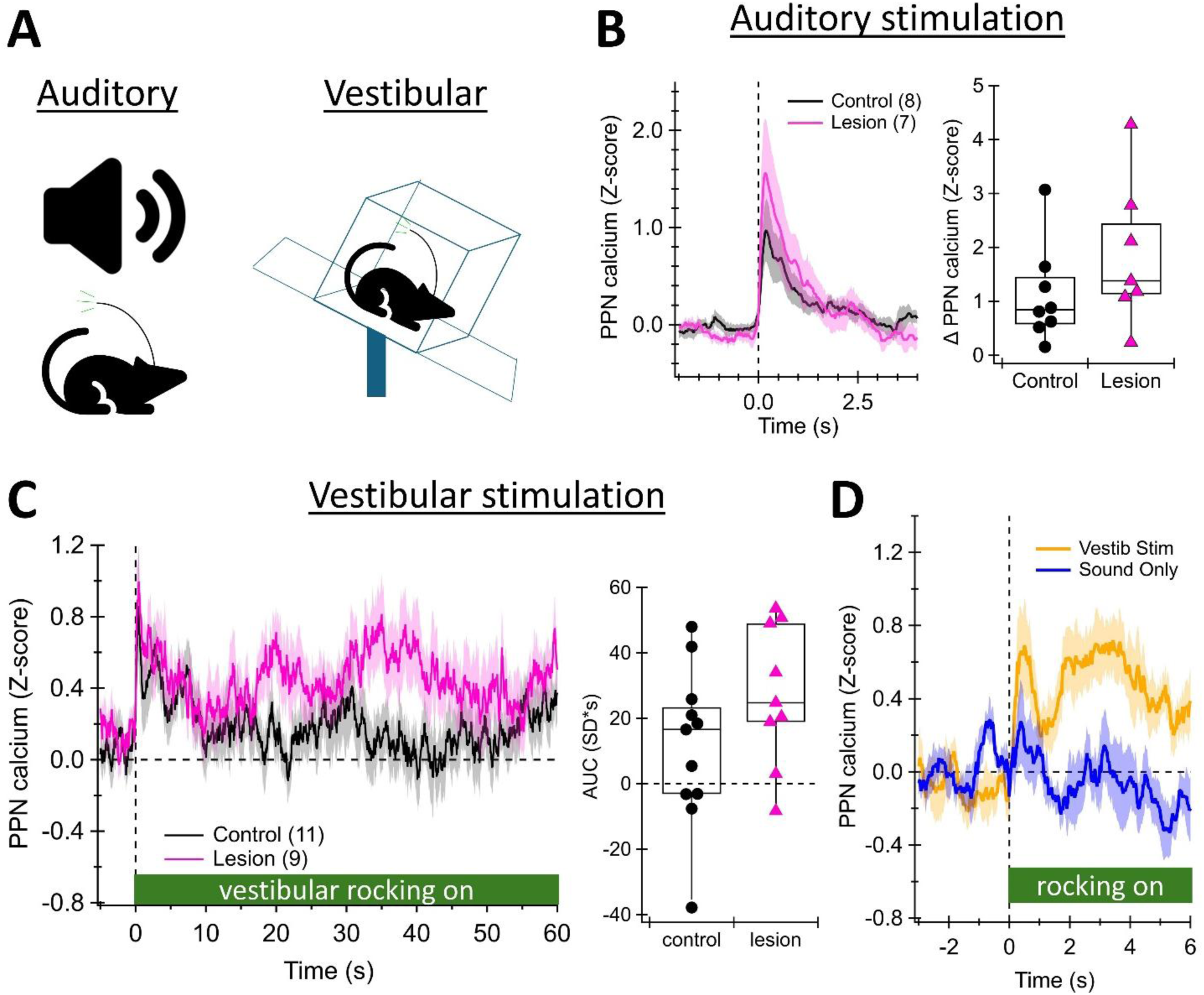
Glutamatergic PPN neurons respond to auditory and vestibular stimuli with and without cholinergic cell loss.. **A.** Schematic of stimuli given during fiber photometry. **B.** Glutamatergic PPN activity in response to auditory tone. Amplitude of response does not differ between control and lesion conditions. **C.** Glutamatergic PPN activity in response to 1 minute of vestibular stimulation. Area under the curve (AUC) of signal does not differ between control and lesion conditions. **D.** Comparison of glutamatergic PPN recordings when the mouse is on the vestibular rocker (Vestib Stim) and when the mouse is next to, but not on the vestibular rocker (Sound Only). Vestibular stimulation results in amplified PPN activity vs the sound of the vestibular rocker.

Because we observed balance deficits in the cholinergic cell loss model (Figure 2A), we hypothesized that cholinergic cell loss would alter the PPN response to vestibular stimulation. To test this hypothesis, we placed the mice in a box on an oscillating rocker and measured PPN glutamatergic population activity during vestibular oscillations when the rocker turned on. In most mice, PPN glutamatergic neurons showed increased calcium activity when the mouse was on the active vestibular rocker as measured by the area under the curve (AUC) during each minute of stimulation (Figure 6C). Because the rocker made sounds when rocking, we ran control stimulation experiments where the mouse was placed in a box next to, but not on, the rocking plate. This generated the same auditory stimulation, but in the absence of vestibular stimulation. Figure 6D shows the average PPN glutamatergic signal from 5 mice recorded on the rocker (yellow) and next to the rocker (blue). This shows that the sound of the rocker alone did not cause a large sustained PPN glutamatergic response. Importantly, there was no significant difference between vestibular response in the lesion vs control group (Lesion 27.34 ± 7.2 SD*s, n = 9; Control 11.4 ± 7.31 SD*s, n = 11; 2-tailed T-test T_(10,8)_ = -1.55, p = 0.14). These data show that cholinergic cell loss does not significantly impact the glutamatergic response to vestibular rocking stimulation.

## Discussion

Here we show that brainstem cholinergic cell loss causes slowly-developing motor symptoms and neural circuit alterations. Importantly, we find physiological changes in glutamatergic PPN neurons, a population previously implicated in motor control ^9,11,13,30,10,31^, after cholinergic cell loss. The physiological alterations include increased excitatory synaptic strength onto PPN glutamatergic neurons and a prolonged glutamatergic response to visual stimuli. These physiological changes occur during the same time frame as the development of motor symptoms.

### Model symptoms relationship to PD and PD models

Several studies in PD patients have shown that within-session motor learning is relatively intact, while motor recall is impaired ^38–40^. In our model of brainstem cholinergic cell loss, we observe this phenomenon, an ability to learn within multiple sequential trials, but a severe loss in learned motor skill after delays of 24 hours or longer. This suggests that the motor recall deficits observed in PD may relate to cholinergic degeneration. Interestingly, motor recall is particularly impaired in PD patients with significant freezing of gait (FOG) ^39,64^. It is possible that PPN degeneration contributes to both FOG development and motor skill recall deficits.

In contrast to the motor recall deficits, we did not find object memory recall impairments using the novel object recognition task, unlike some have found in dopamine-mediated PD models ^65^. These contrasting findings could be explained by the cholinergic PPN having a selective role in motor memory consolidation over spatial over object memory. Alternatively, they could reflect the cholinergic PPN having a selective impact on sleep-related memory consolidation, consistent with previous work implicating the PPN in sleep-wake states ^56,66^. It is important to note that the motor recall deficits were present after a 24-hour delay, while the novel object test was performed after a 1-hour delay, meaning that sleep-related memory consolidation was only a factor in the motor control experiment. Future work will be needed to dissociate these two possible explanations.

Mice with cholinergic cell loss showed balance impairments, consistent with previous work implicating the PPN in postural instability and balance deficits in PD patients and animal models ^4–6,24,44^. Unexpectedly, however, we did not observe significant gait deficits or alterations in this model. This seems to contrast with previous studies implicating the PPN in freezing of gait (FOG) ^67–69^, but is consistent with recent a study showing that extra-PPN cholinergic disruptions are related to FOG development ^70^. Together these data suggest that cholinergic PPN degeneration has a more direct relationship with balance deficits than with FOG.

Mice with brainstem cholinergic cell loss showed an increase in open field locomotion. This finding was surprising because decreased locomotion is a cardinal symptom in PD and many animal models of PD show reduced movement ^71^. However, some progressive animal models of PD show a period of hyperactivity early in disease progression ^34,35,72,73^. The increase in open field locomotion after brainstem cholinergic cell loss could be caused by increases in downstream dopaminergic neural activity or may relate to enhanced PPN glutamatergic neural activity. Indeed, our electrophysiological findings show that PPN glutamatergic neurons receive stronger excitatory synaptic input after cholinergic cell loss.

### Slow development of symptoms

One of the most interesting findings is that motor symptoms did not occur immediately after cholinergic neuron degeneration. Previous studies temporarily inhibiting cholinergic PPN neuron activity show minimal effects on locomotion ^33,74^, but others show decreased locomotion ^75^. Interestingly, temporary chemogenetic inhibition of cholinergic PPN input to the cerebellum actually improves balance ^33^. Together, these findings suggests that secondary circuit adaptations are responsible for balance impairments and increased exploratory locomotion after brainstem cholinergic cell loss. Interestingly, our *ex vivo* electrophysiology experiments show increases in excitatory input to PPN glutamatergic neurons and reductions in intrinsic excitability in response to cholinergic cell loss. Because the glutamatergic PPN population is implicated in motor control ^8,9,11,13^, disrupting circuit connectivity and cellular activity in this population could affect sensorimotor integration.

When recording the PPN glutamatergic response to sensory input, we found robust responses to visual and auditory stimuli, consistent with previous work ^58,76^. We also observe strong sustained PPN glutamatergic activity in response to vestibular stimuli. Because cholinergic cell loss altered PPN glutamatergic neurons at the cellular and synaptic level, we had hypothesized that the sensory response in these neurons would also be impacted. We found a prolonging of the glutamatergic signal in response to visual stimuli at the exact same timepoint as the maximal behavioral effect, 7 weeks post-injection. Although the fiber photometry and behavioral assays were performed in separate groups of mice, the matching timeline suggests an interesting relationship between alterations in sensory processing in the glutamatergic PPN and motor symptom development. Future experiments are needed to determine whether these alterations in the glutamatergic PPN have a causal role in symptom development.

### Causes and consequences of glutamatergic PPN physiological changes

We find increased excitatory input with decreased intrinsic excitability in PPN glutamatergic neurons after cholinergic cell loss. This combination of cellular alterations may reflect increased synaptic potentiation and subsequent homeostatic downregulation of excitability. One possible cause for the synaptic potentiation is that, in the healthy state, cholinergic PPN neurons may tonically inhibit glutamate release onto glutamatergic PPN neurons via pre-synaptic inhibitory muscarinic receptors ^77^. If such tonic inhibition is abolished due to cholinergic cell loss, the excitatory synaptic input may become stronger. Because cholinergic PPN neurons may also directly inhibit PPN glutamatergic neurons in the healthy state ^78^, cholinergic cell loss could increase PPN glutamatergic activity while simultaneously increasing pre-synaptic glutamate release. An increase in both pre-synaptic and post-synaptic activity could result in long-term potentiation of that connection ^79^. While this is a possible mechanism, future experiments will be needed to determine whether local connectivity within the PPN is responsible for these physiological changes after cholinergic cell loss.

Increased synaptic excitation and decreased maximal excitability of glutamatergic PPN neurons could result in one synaptic connection having disproportionate control over glutamatergic neurons. The increase in synaptic current amplitude without an increase in current frequency suggests that new synapses are not necessarily being created, but rather that one or more synaptic inputs are being strengthened. If one input is strengthened, but the neuron becomes less intrinsically excitable, the other synaptic inputs to that neuron will be functionally weaker. Therefore, a major consequence of these physiological changes could be an imbalance in the brain networks controlling PPN glutamatergic output. Because PPN glutamatergic neurons receive excitatory input from many diverse structures in the brain ^8,9^, an imbalance in these inputs could disrupt the motor control computations performed by the PPN.

### Conclusions

In this study, we selectively lesioned cholinergic brainstem neurons to isolate the symptoms and circuit alterations that occur. We found that cholinergic cell loss causes cellular and circuit alterations in the brainstem that accompany slowly-developing motor symptoms. Cholinergic cell loss did not affect any non-motor symptom that we tested. These findings provide causal insight into the relationship between brainstem cholinergic cell loss and PD motor and non-motor symptoms.

## Methods

### Animal Subjects

All animal procedures were approved by the Georgetown University Medical Center Institutional Animal Care and Use Committee (IACUC). Measures were taken to ensure minimal animal suffering and discomfort, and protocols were designed to minimize the number of animals used. Homozygous Ai9-TdTomato mice [B6.Cg-Gt(ROSA)26Sortm9(CAG-tdTomato)Hze/J; JAX #007909] were bred in-house with heterozygous ChAT-Cre mice [B6.129S-Chattm1(cre)Lowl/MwarJ; JAX #031661] to obtain ChAT-cre heterozygous/Ai9 heterozygous and Ai9 heterozygous (Wild-type littermates). Experiments were conducted using mice at least 8 weeks old of both sexes. Animals were housed under a 12:12 h light– dark cycle (lights on at 06:00 AM) with food and water available *ad libitum*.

### Surgical Procedures

Mice were anesthetized with 5% isoflurane in an induction chamber and then transferred to a stereotaxic frame (Stoelting 51730UD). Anesthesia was maintained with 1-3% isoflurane delivered in oxygen at a constant flow rate of 1 L/min. Body temperature was maintained using a warm pad. The skull was stabilized using ear bars. To avoid damage to the ears which might impair performance in vestibular and auditory tasks, ear bars were placed in the zygomatic arch rather than in the ear canal. Bupivacaine (5 mg/kg) was administered subcutaneously at the incision site as a local anesthetic, and mice received carprofen (5 mg/kg) as a general analgesic prior to surgery. A small scalp incision was made to expose the skull, and bregma and lambda were identified to ensure proper leveling of the head.

### Viral Injections and Fiber Implantation

Bilateral craniotomies were drilled above the PPN and intracranial injections were given at coordinates AP −4.5 to -4.7 mm, ML ±1.1 mm, DV −3.7 mm relative to bregma. A 33-gauge needle was lowered to the target, and 1 µL total volume was injected with a 5-µL Hamilton micro-syringe at a rate of 0.2 µL/min. The syringe was held in place for 1 minute following the injection, then raised 0.5 mm above the target site and left in position for an additional 10 minutes to minimize backflow. For behavioral experiments, AAV5-FLEX-taCasp3-TEVp (1.6 × 10^13^ particles/mL) or AAV5-hSyn-DIO-EGFP (1.1 ×10^13^ particles/mL) was injected as lesion or control virus, respectively. For electrophysiology of opto-tagged PPN glutamatergic neurons, AAV-CamKIIa-ChR2 (1 ×10^13^ particles/mL) was mixed (1:1 to 1:4) with the AAV5-FLEX-taCasp3-TEVp virus or sterile PBS and 500nl per hemisphere injected bilaterally into the PPN of ChAT-cre/Ai9tdTomato mice. For fiber photometry experiments, AAV-CamKIIa-jGCaMP8f-WPRE (2.0 ×10^13^ particles/mL) was mixed (1:1) with AAV5-FLEX-taCasp3-TEVp or sterile PBS (1:1) and injected bilaterally into the PPN of ChAT-Cre/Ai9-tdTomato mice. An optical fiber (400-µm core, 0.22 NA, 3.7-mm length) was additionally implanted in the left hemisphere above the PPN and secured using dental cement (C&B-Metabond Quick! and Jet Denture Repair). For behavioral experiments, the incision was closed with Surgical Sutures (Stoelting Visorb Polyglycolic Acid (PGA) Synthetic Absorbable Sutures) or wound clips. Postoperatively, mice received either carprofen, buprenorphine SR or both and recovered for a week prior to the start of post-lesion behavioral testing.

### Behavioral Experimental Groups

Mice were assigned to one of three groups: the Lesion group, consisting of ChAT-Cre/Ai9-tdTomato mice that received AAV5-FLEX-taCasp3-TEVp to selectively ablate PPN cholinergic neurons (n = 16; 7 females, 9 males); the ChAT-Cre Control group, consisting of ChAT-Cre/Ai9-tdTomato mice that received AAV5-hSyn-DIO-EGFP (n = 17; 8 females, 9 males); and the WT Control group, consisting of Wild-type littermates (with the exception of 7/20 WT mice which were age-matched but not littermates) that received AAV5-FLEX-taCasp3-TEVp as a virus control (n = 20; 9 females, 11 males).

### Behavioral Tasks

With the exception of voluntary wheel running which took place in home cages over a period 7 days, all behavioral assessments were conducted during the light phase, and testing chambers were cleaned with 70% ethanol between mice to minimize olfactory cues. Mice were left in the behavior room for about 30 minutes before testing to acclimate the mice to the testing environment.

#### Open Field

To evaluate longitudinal locomotor activity and anxiety-like behavior, mice were subjected to the open-field test (OF) and an elevated zero-maze (EZM) before and after viral injections. Subject locations and exploratory behaviors across both paradigms were continuously monitored and quantified using ANY-maze video tracking software (version 7.8). For the OF test, mice were placed in the center of a square arena (45 × 45 cm, illuminated at ∼400 lux) and allowed to freely explore for 10 min. Baseline behavior was established prior to viral injection, followed by weekly testing for eight consecutive weeks. Quantified metrics included total distance traveled, movement speed, and time spent in the center versus as well as the relative distance traveled and entry frequency into the center zone (a 22.5 × 22.5 cm square).

#### Balance Beam Task

Motor coordination and balance were assessed by averaging the time taken to cross beams of three widths (30mm, 6mm, and 3mm) elevated 50 cm above the bench surface. Mice were trained to traverse to a darkened escape box (120 cm away). Prior to stereotaxic surgery, mice underwent five consecutive days of training. On each day, mice crossed the 30 mm beam twice, the 6 mm beam three times, and the 3 mm beam three times. Each mouse was given a maximum of 120 seconds to traverse the beam per trial. Mice that were unable to cross the 3 mm beam in under 120 seconds on Day 5 were excluded from balance beam analysis. After stereotaxic surgery, mice were tested once per week following the same procedure (save one cohort which was run twice per week). The time required to cross the beam was recorded and the trials per width were averaged. Averages for time to cross across all cohorts were grouped and averaged based on their respective lesion condition and sex differences. Data was normalized to the average time to cross on the last (5^th^) training day prior to surgery.

#### Novel Object Recognition

Two novel object recognition sessions were conducted: one 5 days prior to lesion surgery and a second session 6 weeks after. All procedures were conducted in a 45 × 45 cm open-field arena (400 Lux). The weekly open-field session served as Day 1 chamber habituation. On Day 2, mice were placed in the arena with non-test objects for 10 minutes. On Day 3 (test day), identical objects were placed in opposite corners. In the familiarization phase, mice were allowed to explore freely for 10 minutes; trials were included only if mice accumulated at least 30 seconds of total exploration time. After a 1-hour delay, one familiar object was replaced with a novel object for the test (retention) phase. Object recognition memory was quantified using a discrimination index: (time spent exploring novel – time spent exploring familiar) / total exploration time.

#### Gait Analysis

Gait assessment was performed using the CatWalk XT Gait Analysis System (version 10.1). A high-speed camera recorded the position (x, y) and timing (t) of each paw contact at a self-selected walking speed. For each experimental time point, the gait model from that week was compared to the baseline model to compute a multidimensional locomotor error metric. The relative change in locomotor error across weeks was used as a measure of overall locomotor impairment or recovery ^45^.

#### Elevated Zero Maze

Anxiety-like behavior was assessed using a circular elevated maze (65 cm diameter, 5 cm wide track, 51 cm height) divided into two open and two enclosed quadrants. Mice were placed in an open quadrant and allowed to freely explore for 10 minutes. Primary measures included time spent in open versus closed quadrants and number of entries into open arms. Post-lesion testing was conducted on Weeks 4, 6, and 10. Time spent in each quadrant was recorded. Data was normalized as a percentage of the time spent in the open arms during pre-surgery assay.

#### Prepulse Inhibition (PPI)

Sensorimotor gating was assessed using the SR-LAB system. The session consisted of a 70 dB background noise presented alone for 5 minutes as acclimation. Following acclimation, mice were presented with five “Pulse Alone” trials (120 dB, 30 ms) for habituation. These were excluded from analysis. After habituation, mice were presented with “Pulse Alone” and “Prepulse+Pulse” stimulation events, with prepulses 3, 6, 9, and 12 dB above background. If startle amplitudes after PPI assays were lower than 40% of baseline startle amplitude, data were excluded. Each stimulation type was applied 5 times total and stimulation types were presented pseudo-randomly with varying inter-trial-intervals. Prepulses were given 130 ms before the accompanying pulse. PPI was calculated as: % PPI = 100*[1 − (startle response for prepulse + pulse /startle response for pulse-alone)]. Mice were tested prior to lesion induction and post-lesion on Weeks 4 and 8.

#### Rotarod

Motor coordination and learning were assessed using the Ugo Basile Rotarod (47650). On week 10 post-lesion, mice underwent four consecutive days of training (three trials per day) and then a final test day six days after the fourth training day. The rod accelerated from 5 to 80 rpm over six minutes. Inter-trial intervals were 10–15 minutes. Latency to fail was recorded for each trial. Failure occurred when the mouse fell off the rod or clung to the rod and circled with it two times in a row. To specifically evaluate overnight motor retention, a “rotarod recall” score was calculated for each mouse. This metric was defined as the difference in latency to fail between the first trial of a given day and the final trial of the preceding day, averaged across all training days.

#### Voluntary Wheel Running in Cage

Following behavioral tests in Week 12, mice were single-housed and received a low-profile wireless saucer-style running wheel (MedAssociates) in their home cage. Mice had unlimited access to the wheel for 7 days, and voluntary running activity was continuously recorded using magnetic sensors to track wheel revolutions. Activity was recorded and tracked using MedAssociates Wheel Manager Data Acquisition Software.

### Ex vivo whole-cell electrophysiology

Mice were randomly assigned to one of two groups: the Lesion group, consisting of ChAT-cre/Ai9-tdTomato mice that received AAV5-FLEX-taCasp3-TEVp to selectively ablate PPN cholinergic neurons (n = 11; 4 females, 7 males) and the Control group, consisting of ChAT-cre/Ai9-tdTomato mice that received sterile PBS (n = 9; 4 females, 5 males). Both groups also received AAV-CaMKIIa-ChR2 with this injection to optically tag PPN glutamatergic neurons for whole-cell recording. 1–8 glutamatergic neurons were recorded per mouse.

5–8 weeks after viral injection, mice were anesthetized with isoflurane and transcardially perfused with cold, oxygenated slicing solution (in mM: 198 glycerol, 2.5 KCl, 1.2 NaH_2_PO_4_, 20 HEPES, 25 NaHCO_3_, 10 glucose, 10 MgCl_2_, 0.5 CaCl_2_). Brains were extracted into cold slicing solution and 200 µm-thick coronal slices of the PPN were obtained using a VT1000S Leica vibratome. The slices were incubated in 34°C oxygenated holding solution (in mM; 92 NaCl, 2.5 KCl, 1.2 NaH_2_PO_4_, 30 NaHCO_3_, 20 HEPES, 2 MgCl_2_, 35 glucose, 5 sodium ascorbate, 3 sodium pyruvate, 2 thiourea) for 30 minutes and then at room temperature for at least 30 minutes. During recordings, slices were superfused in a recording chamber with oxygenated artificial cerebrospinal fluid (in mM: 125 NaCl, 25 NaHCO_3_, 3.5 KCl, 1.25 NaH_2_PO_4_, 10 glucose, 1 MgCl_2_, 2 CaCl_2_) at a temperature of 30–34°C. Each of these solutions had an osmolarity of 300–310 mOsm.

Electrophysiological signals were recorded using an Axon Digidata 1550B digitizer and a Multiclamp 700B amplifier. Molecular Devices Clampex software version 11.2 was used for acquisition. Both voltage clamp and current clamp recordings were sampled at 10 kHz. Voltage clamp recordings were low-pass filtered at 2kHz, while current clamp recordings were low-pass filtered at 10 kHz. Slice tissue was visualized using an Olympus OpenStand upright microscope and light was delivered by ThorLabs LED.

Borosilicate glass micropipettes of 1.5–5 MΩ resistance were made using a Sutter Instrument P-97 pipette puller and filled with internal solution (in mM: 122 KMeSO_3_, 9 NaCl, 9 HEPES, 1.8 MgCl_2_, 14 phosphocreatine, 4 Mg-ATP, 0.3 Tris-GTP, 0.09 CaCl_2_, 0.45 EGTA) with 0.08-0.09% w/v neurobiotin. with an osmolarity of 290–300 mOsm and pH of ∼7.35. Cholinergic neurons expressing Ai9-tdTomato were identified using 565 nm light and non-fluorescent (non-cholinergic) cells in the PPN were targeted for recording. The ChR2-expressing nature of recorded cells was verified in voltage clamp using a two-second period of 470 nm light while holding the cell at -70 mV. Cells tagged with the ChR2 exhibited a sustained inward current throughout the light stimulation. Cells that did not exhibit this current were considered putative GABAergic neurons and were excluded from analysis.

#### Spontaneous Excitatory Input

To record spontaneous excitatory postsynaptic currents (sEPSCs), 200 seconds were recorded in voltage clamp while holding cells at -70 mV. Any cells that had an access resistance greater than 25 MΩ to hold at -70 mV were excluded from analysis. Analysis of sEPSC characteristics including amplitude, frequency, decay tau, and 10–90% rise time was performed using MiniML software ^80^.

#### Spontaneous Firing

To measure spontaneous firing activity, 20 seconds were recorded in current clamp in gap-free mode with no holding current applied. The frequency of action potentials was analyzed using custom scripts on Igor Pro. Cells that required hyperpolarizing current to fire spontaneously were excluded from analysis.

#### Intrinsic Excitability

To measure intrinsic excitability and rebound parameters, a current-voltage curve was recorded in current clamp using a series of one-second hyperpolarizing and depolarizing current injections starting at -100 pA and ascending in increments of 25 pA to a maximum of 175 pA. If the cell entered depolarization block during current injection, the recording was stopped. For the current-evoked spiking analysis, any cells that required a hyperpolarizing holding current greater than -80 pA to fire spontaneously were excluded from analysis. Measures such as maximum action potential frequency during depolarization, firing adaptation index during depolarization, and rebound activity after hyperpolarization were extracted using custom scripts on Igor Pro.

### Fiber photometry

Continuous dual-wavelength fiber photometric recordings were measured using the Neurophotometrics FP3002 system. A low-autofluorescence fiber optic patch cord (400-µm core, 0.37 NA, Doric) was attached to the implanted optic cannulas using 1.25mm inner diameter mating sleeves. The PPN was illuminated with 470 nm excitation light to excite GCaMP8f fluorescence, while a 415 nm isosbestic control channel was utilized to account for motion artifacts and photo-bleaching. Light power for both the 470 nm and 415 nm channel were held at 6% of maximum output and power was measured at ∼125-130 µW, measured at the patch cord tip at constant illumination using a Thor labs PM20A optical power meter. All data acquisition and behavioral synchronization were managed using custom workflows in the Bonsai open-source visual programming framework ^81^. Recordings were made at 30 Hz per channel.

### Sensory Stimulation Paradigms

After stereotaxic surgery, mice were held for a 3-week post-surgery period to allow for full protein expression of the GCAMP8f and caspase viruses. Following this 3-week period, visual, auditory, and vestibular sensory modalities were assessed in lesioned and control mice.

#### Visual Stimulation

To assess state-dependent circuit changes, we tested the glutamatergic neuronal response within the cholinergic-lesioned PPN to acute visual stimuli. Ambient lighting intensity within the testing arena was controlled via an Arduino-based microcontroller, allowing for precise transitions between 137 lux (light ON) and 0 lux (light OFF). Mice were first subjected to a 5-minute acclimation period under standard room lighting conditions (137 lux). Following this baseline, the ambient light was turned OFF and ON 4 times. To prevent habituation, the duration of these stimulus epochs varied pseudo-randomly between 50 and 62 seconds (Light OFF durations: 58, 52, 60, and 59 seconds; Light ON durations: 52, 62, 50, and 50 seconds).

#### Auditory Stimulation

To evaluate auditory responsiveness, testing was conducted at 6 and 9 weeks post-lesion within a standard 40 × 40 cm open field arena. Trials commenced with a 180-second acclimation baseline in the presence of continuous 70 dB ambient white noise. Acoustic stimulation was subsequently generated via two 3–24V active piezo buzzers configured to emit a 500 ms 82 dB, 2800 - 3200 Hz tone. The tone was delivered four times throughout the session 90 seconds apart

#### Vestibular Stimulation

Vestibular testing was conducted concurrently with the auditory assessments at 6 and 9 weeks post-lesion. Mice were placed within a 10 × 10 cm acrylic enclosure mounted to a 30 cm² Blot Boy platform shaker. To induce vestibular activation, the shaker was operated in a manually timed ON/OFF sequence. The active stimulation consisted of a continuous, low-frequency oscillation cycle where the shaker platform moved through a vertical displacement of 6 cm over 5 seconds before returning to the starting position. Given that the shaker apparatus produces a distinct mechanical noise during oscillation, a dedicated sound control was implemented to parse vestibular-specific neural signals from auditory artifacts. During these control sessions, the testing chamber was positioned immediately adjacent to the active shaker, exposing the mice to the acoustic profile and timing intervals without the physical motion. Stimulus onset and offset for both the active motion and sound-control trials were time-stamped via a keydown event in the Bonsai workflow.

### Fiber Photometry Analysis

Raw fluorescence intensity measurements from the 470 nm (calcium-dependent) and 415 nm (isosbestic reference) channels were acquired continuously via the Bonsai system. To parse true calcium-dependent neural activity from motion artifacts and photobleaching, the 415 nm reference signal was subtracted from the 470 nm GCaMP8 signal and z-scored.

### Brain Perfusion, Slicing and Clearing

At the conclusion of experiments, mice were deeply anesthetized and transcardially perfused with phosphate buffer (PB) followed by 4% paraformaldehyde (PFA). Coronal brain sections (200-μm thickness) were obtained using a vibratome (PELCO easi-Slicer) and collected in PB. Tissue sections were then processed using a CUBIC tissue clearing protocol ^32^. To facilitate clearing, sections were incubated in CUBIC Reagent 1 (R1) overnight at room temperature on an orbital shaker. Sections from caspase-lesioned and ChAT control animals were subsequently transferred directly to CUBIC Reagent 2 (R2) prior to mounting on slides. All incubation steps, including CUBIC R1 delipidation, antibody labeling, and CUBIC R2 refractive index matching, were performed at room temperature with continuous agitation.

### Confocal Microscopy and Image Processing

Fluorescence images were acquired using a laser scanning confocal microscope (SP8, Leica) equipped with a 10× objective. Z-stacks were acquired with 5-micron steps and 20 slices total. To quantify cell density, confocal Z-stacks were processed using a custom automated macro in ImageJ/Fiji. Multi-channel stacks were split to isolate the specific fluorescence signal of interest. To generate maximum intensity projections, Z-stacks were divided into 4 consecutive 5-slice substacks. Prior to image segmentation, a despeckle filter was applied to reduce background noise. Images were then binarized using the MaxEntropy thresholding algorithm (dark background), followed by a watershed transformation to separate adjacent or clustered cells. Finally, automated particle analysis was performed to quantify cell numbers in each of the 4 projected substacks. Inclusion criteria were restricted to objects with a minimum area of 10 pixels and a circularity index of 0.50–1.00. Cell counts were summed for each substack of each hemisphere of each mouse. Hemispheres were averaged together to get the total cell count per slice per mouse (Figure 1).

### Statistical Analysis

All behavioral and histological data were analyzed using SAS (version 9.4) or Igor Pro (version 9.0). Data are presented as mean ± SEM unless otherwise indicated. Appropriate statistical tests, including ANOVA, repeated-measures mixed-effects models, and post hoc tests with corrections for multiple comparisons were applied. Shapiro-Wilk tests were used to assess normality of the data. If data were normally distributed, ANOVA and t-tests were applied. If data were non-normally distributed, Wilcoxon Mann-Whitney U tests were applied to compare two conditions. Statistical significance was defined as p < 0.05.

## Acknowledgements

This work was supported by the Michael J Fox Foundation #MJFF-022863, a Parkinson’s Foundation Stanley Fahn Junior Faculty Award #PF-SF-JFA-1040267, and NINDS #R00NS112417 awarded to RCE, an NINDS supplement #R00NS112417-S1 awarded to RCE and MEO, by an NIH NINDS grant #1F30NS132399 awarded to MF, a Parkinson’s Foundation Summer Student Fellowship #PF-SSF-1646817 awarded to GZ, NIH NINDS T32NS041218 awarded to BJB, and NIH NIA T32AG071745 awarded to VJL. We thank the Georgetown University Microscopy and Imaging Shared Resource (MISR) for use of the confocal microscope and the Neuroscience Department for use of the animal behavior core facilities. We also thank members of the Evans lab for feedback on versions of this manuscript.

## Notes

### Competing Interest Statement

The authors have declared no competing interest.

